# Building Bridges: Mycelium Mediated Plant-Plant Electrophysiological Communication

**DOI:** 10.1101/2022.07.20.500447

**Authors:** Matthew Adam Thomas, Robin Lewis Cooper

## Abstract

Whether through root secretions or by emitting volatile organic compounds, plant communication has been well-documented. While electrical activity has been documented in plants and mycorrhizal bodies on the individual and ramet, electrical propagation as a means of communication *between* plants has been hypothesized but understudied. This study aimed to test the hypothesis that plants can communicate with one another *electrically* via conductively isolated mycelial pathways. We created a bio-electric circuit linking two plants using a mycelial network with a blend of endomycorrhizal fungi grown on potato dextrose agar forming the isolated conductive pathway between plants. Using this plant-fungal biocircuit we assessed electrical propagation between *Pisum sativum* and *Cucumis sativus* We found that electrical signals were reliably conducted across the mycelial bridges from one plant to another upon the induction of a wound response. Our findings provide evidence that mechanical input can be communicated between plant species and opens the door to testing how this information can affect plant and fungal physiology.

**Simple Summary:** Most plants form underground relationships with fungi. These relationships are mutually beneficial. The plants and fungi share, trade, and distribute resources between themselves, their neighbors, and their offspring. Plants employ diverse methods to detect and respond to their environment and the production of electric signals is one of these methods. It would be favorable to a plant’s survival and the survival of their neighbors, if this plant could transmit and share the information these electrical signals contain. Possible avenues of transmission exist in the roots, and the fungi these roots are in contact with. If a fungal mass is in contact with the roots of multiple plants, it could propagate electrical signals throughout the plant network. We found that electric signals were reliably transmitted from one plant to another via fungal pathways upon the induction of a wound response. Our findings provide evidence that mechanical input can be communicated between plant species and opens the door to testing how this information can affect plant physiology.

## 1. Introduction

“The fact that we lack the language skills to communicate with nature does not impugn the concept that nature is intelligent, it speaks to our inadequacy for communication.”
-Paul Stamets, *Fantastic Fungi*

Plant electrophysiology is an exciting and widely expanding field. Scientists are developing new methods for signal detection (intracellular and extracellular), writing new software for classifying the diverse signal types plants produce [1], and studying how various signals regulate genetic expression which induce a wide array of complex responses. Fungal electrophysiology is also gaining traction as a new frontier linking large biotic plant communities. As the scientific community delves deeper into these evolving fields, one is beginning to understand just how complex electrical communication networks among organisms may be.

Roughly 90% of all plants form symbiotic relationships with mycorrhizae [2]. Experiments have shown that common mycelial networks can warn neighboring plants of aphid attack [3] and facilitate interplant nutrient exchange [4]. The mechanisms underlying these interactions are complex and require specific *in situ* and *in vitro* methodologies for deciphering these interactions.

Arbuscular endomycorrhizal fungi (AMFs) penetrate cell walls but not cell membranes in plant roots and form structures called arbuscules. Since the fungi penetrates intracellularly, it gains direct and indirect access to the symplast [5]. Although the arbuscule penetrates the cell wall, it never enters the host’s cytoplasm directly, but instead interfaces with the cytoplasm through a three-layered structure called the periarbuscular membrane (PAM). The PAM acts like a multimolecular trading post which promotes active and bidirectional transport of nutrients and is capable of producing energy gradients [6]. Where there are energy gradients and ionic differences, there is an electric potential which can produce changes in voltage.

Both plant and fungal electrophysiology has been studied in isolation, but studying electrical interactions between phylogenetically distant organisms has proven difficult because this communication occurs almost exclusively underground. The conductivity of soil varies based on ionic composition and moisture, but in general, soil is a conductor of electricity. Furthermore, the agar used for culturing mycelium in the lab is itself conductive - making it difficult to pinpoint whether the mycelium, or the growth medium, conducted an induced electrical signal over a given distance. Even if one could directly observe the electrical interactions occurring in the soil between roots and fungi it is challenging to definitively determine if the mycelium conducted a signal as opposed to the soil around the symbionts.

We report a method for studying mycelium-mediated plant-plant electrophysiological communication by means of isolating the conductive pathway of electrical responses that can direct the plant signals across mycelial bridges from one plant to another.

## 2. Materials and Methods

### 2.1. The Mycelial Bridge Setup - Agar Inoculation with no host present

Petri dishes with Potato Dextrose Agar (1% w/v) from Seaweed Solution Laboratories (Trondheim, Norway) were used as the growth medium for mycelium. Although endomycorrhizae are obligate endosymbionts requiring a host to survive, studies have shown that various endo and ectomycorrhizal fungi can be grown in culture on potato dextrose agar from propagules [7,8]. Since these are not the most ideal growing conditions for endo/ectomycorrhizal fungi, further studies were conducted which directly inoculated the roots and seeds of the plants being tested.

The Petri dish and agar were cut in half. The agar on each half of the Petri dish was then cut again to have a trapezoidal face with a rounded base (Figure 1A). Both halves of the agar design are denoted with the title “agar islands.” This agar geometry encouraged the mycelium to bridge the gap in the center of the setup - as this was the nutrient path of least resistance. Once the agar was cut to the proper geometry, it was then inoculated with MycoGrow for Vegetables from Fungi Perfecti (Olympia, WA, USA) (286,598 Propagules per kg Total) *Glomus intraradices*, *Glomus aggregatum*, *Glomus mosseae*, *Glomus etunicatum* (71.7 propagules/g each). The method of inoculation was a light powdering over the surface of each agar island - the inoculant can be seen on the agar in Figure 1C. The two halves of the dish were fastened together while maintaining a gap between the two islands of agar. This ensured there was no conductive pathway between the two islands. This gap, as well as the methodology used to cut the Petri dishes in half, made it impossible to maintain sterility throughout the experiment. Immediately after cutting the dishes, the agar was inoculated with MycoGrow for Vegetables. The authors would like to note that although this endomycorrhizal blend was inoculated in the absence of a host, mycelium grew. The authors cannot rule out contamination at this stage of experimentation, but importantly, fine filamentous structures nevertheless grew on the dish which visually resembled mycelium. Furthermore, the organisms which grew bridges of mycelium-like structures looked very similar to the bridges formed in the latter trials in which the host plant was directly inoculated with inoculant.

**Figure 1.**
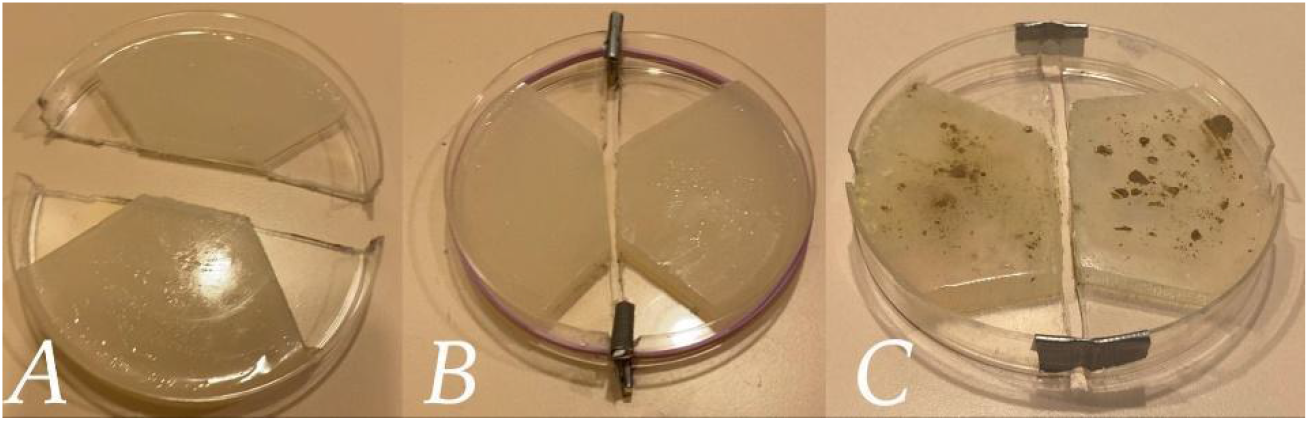
Petri Dish Design. Figure 1A illustrates the Petri dish cut in half with the agar on each side cut into the design to guide the mycelium across the agar gap in the center of the dish. Figure 1B depicts the Petri dish reassembled with spacers and rubber bands in place. Figure 1C represents the second method for reassembling the Petri dish using duct tape. Figure 1C also illustrates how the MycoGrow inoculant appeared on the agar after application.

Two methods of fastening the Petri dishes were used - each method provided its own set of challenges. The first method involved separating the two halves of the dish with small wooden spacers at the edges, and then using a tensioning rubber band to keep the two halves of the dish securely together (Figure 1B). This method proved to be difficult to work with because the tension from the rubber band could make the setup unstable if held or moved in the wrong way. The second method for fastening the two halves of the dish together was much simpler, but came with a different set of challenges. This method involved simply taping the two halves of the dish together at the edges with tape - being conscious not to create a conductive pathway between the two islands of agar while maintaining the necessary gap between islands. Although not as rigidly secure as the method with the wooden shims and rubber band, the tape method required less assembly time. One can observe in (Figure 1C) this method of arranging the agar islands together as well as how the MycoGrow was applied to the agar. Since the dishes were incubated between 18-21 C in a humidity chamber, the adhesive properties of the tape weakened and required re-adhesion every 3-5 days.

The humidity chamber was a plastic container with a slightly damp paper towel placed inside. This extra moisture seemed to greatly increase the inoculation success rate, but more importantly, the number of bridges which formed. The dishes were allowed to incubate and grow for at least two weeks. After incubation, each dish was visually inspected for mycelial bridges, and the best bridgings were chosen for testing. Examples of these bridgings are illustrated in (Figure 2).

**Figure 2.**
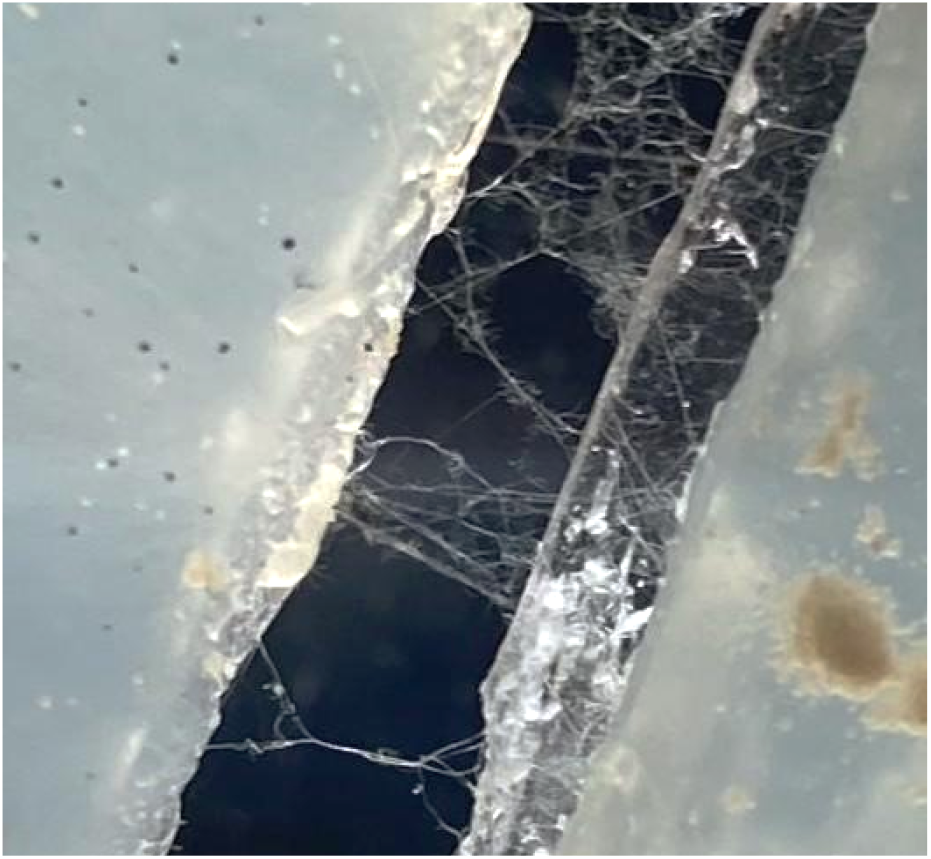
Mycelial bridging across the two islands of agar.

The agar of mycelial bridges were placed on a testing platform which mimicked the gapping of the agar dish itself, but to a greater degree (Figure 3). This gap ensured no conductive interference from underneath the setup. A sheet of paper, either red or black, was placed under the testing platform. This colored paper aided in the visual confirmation of mycelial bridges during testing by providing a contrast to the white mycelial threads. An example of this contrasting background can be seen in Figure 2. The methods described in 2.1 were implemented in Pea Trials 1 and 2 and Cucumber Trials 1 and 2.

**Figure 3.**
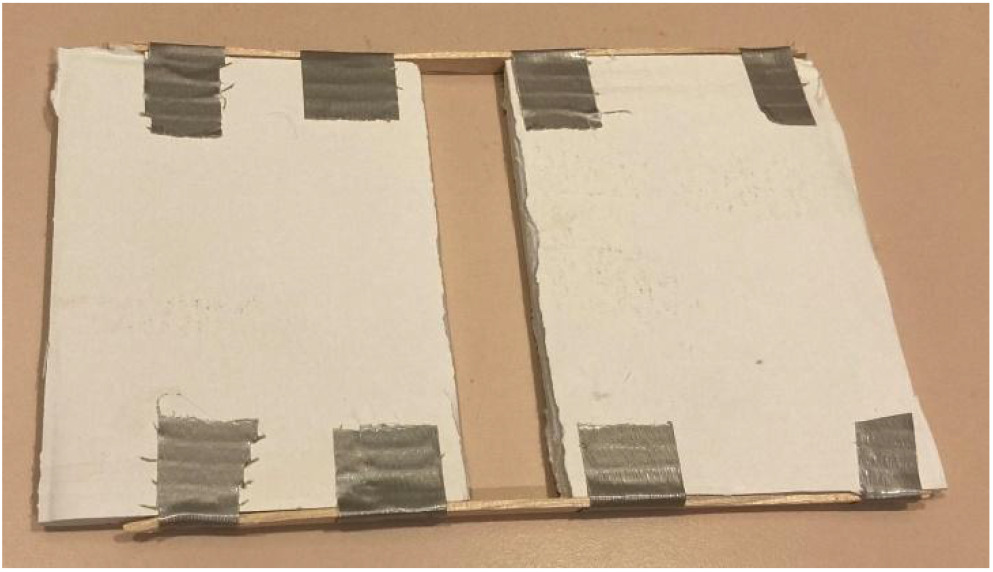
Setup Testing Platform. This figure illustrates the testing platform used to aid in the isolation of the signal. The gap in the center of the platform kept the Petri dish off the surface of the table. This ensured the dish did not touch the testing table and further suspended the mycelial bridge in midair.

### 2.2 The Mycelial Bridge Setup - Direct Inoculation with host/symbiont present on agar

Due to the symbiotic nature of endo/ectomycorrhizal fungi, a different method of inoculation was used to increase the efficiency and efficacy of the mycelial bridges following the first four trials. Due to the methodology presented, contamination is impossible to avoid, but measures can be taken to give the fungi trying to be grown an advantage. Starting with agar which had been autoclaved and poured into Petri dishes under a fume hood helps keep the growth medium sterile until ready to be used. *P. sativum* was directly inoculated with MycoGrow: Micronized endo/ectomycorrhizal fungi sold by Fungi Perfecti (Olympia, WA, USA) (286,598 Propagules per kg Total) *Glomus intraradices*, *Glomus aggregatum*, *Glomus mosseae*, *Glomus etunicatum* (71.7 propagules/g each), *Rhizoogon villosullus*, *R. loteolus*, *R. amylopogon*, *R. fulvigleba* (2,750 propagules/g each), *Pisolithus tinctorius* (220,509 propagules/g), *Scleroderma cepa* and *S. citrinum* (5,500 propagules/gm) to broaden the types of species present in the medium and to increase the chances of successful inoculation. By directly inoculating the pea’s roots, or the pea seed shortly after germination, you give the endo/ectomycorrhizal fungi a head start against possible airborne contaminants that might land on the agar. When using pea seeds, the seed was placed in a bag of inoculant and shaken until coated thoroughly. The inoculant-coated seeds were then placed on each island of agar respectively (Figure 4A) and a damp paper towel was placed over them to retain moisture (Figure 4B). When inoculating pea plants with well established root systems, .05 grams of inoculant was powdered over the root system which rests on each island of agar respectively (Figure 4C), then a damp paper towel was placed over the root systems of both plants to retain moisture for the sake of keeping the roots healthy (Figure 4D). The lids of the dishes were put back on to help retain moisture. For the larger pea plants, small holes need to be cut in the lids to allow the stems to grow upwards. The dishes were then secured to the testing platform to stabilize the setup. Finally, the setups were placed in large plastic containers at around 21° Celsius and incubated for approximately two weeks before testing. All setups were examined under a microscope prior to testing and visual confirmation of bridges were acquired for all setups. An example of a successful bridging when viewed under the microscope can be seen in Figure 5. The methods described in 2.2 were used for Pea Trial 3.

**Figure 4.**
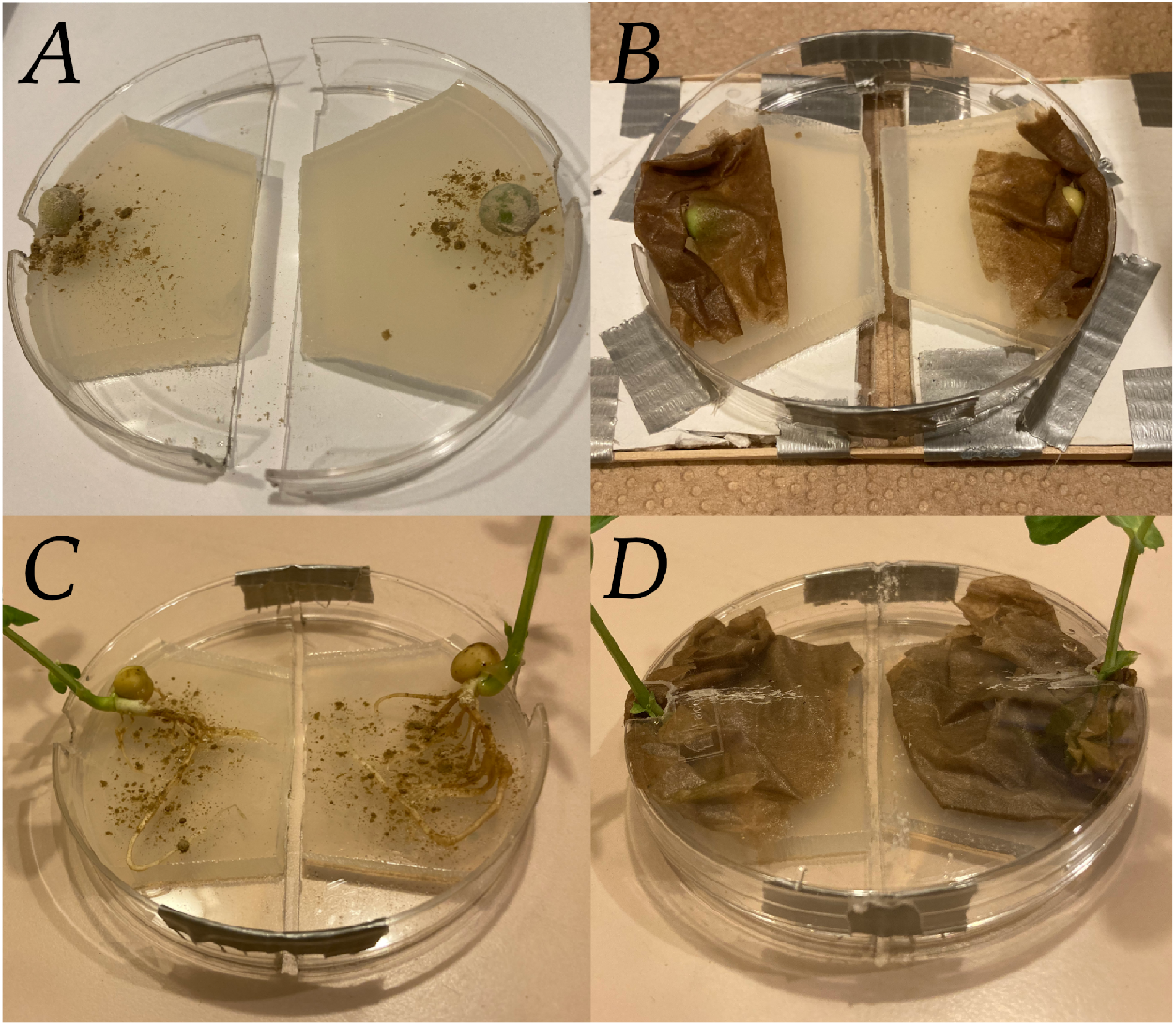
The Mycelial Bridge Setups with direct inoculation of pea seeds and developed pea roots. Figure 4A depicts the pea seeds just after being coated with the Endo/Ecto MycoGrow Inoculant. Figure 4B depicts the same pea seeds, now covered with a damp paper towel to maintain moisture, the dish is now fastened together with tape at the ends while maintaining the necessary gap. The dish has also been fastened to the testing platform for stabilization. Figure 4C depicts direct inoculation of the pea’s root systems. Figure 4D depicts these same pea plants with the roots now covered by a damp paper towel to maintain moisture. This setup would also have been fastened to a testing platform, as seen in 4B.

**Figure 5.**
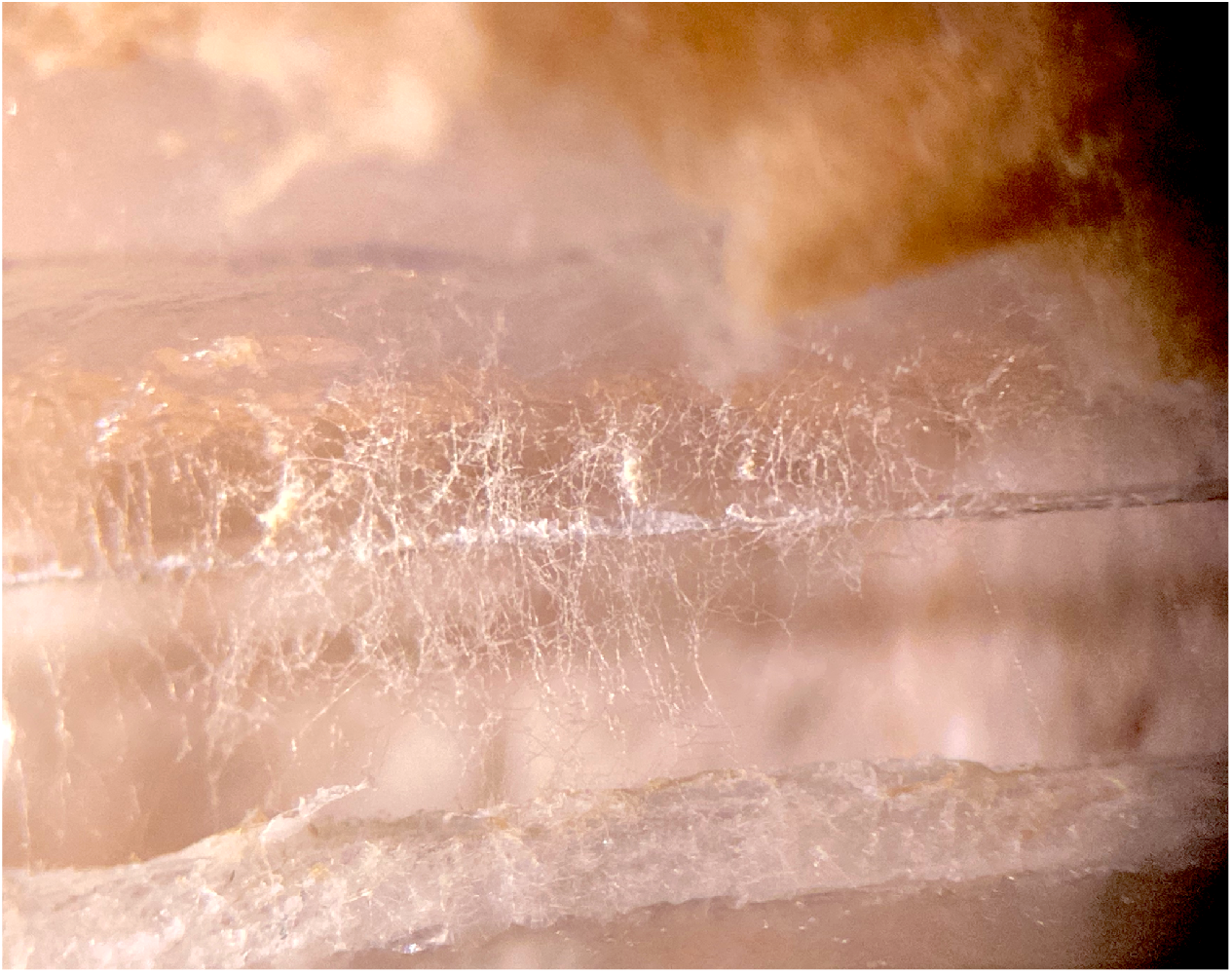
Mycelial bridge when viewed under a microscope. This figure depicts the presence of a mycelial bridge which the authors would visually confirm under the microscope prior to testing the setup.

### 2.3. Growing the Plants

One thing my pea plants taught me: always do science with things you can make into soup.
- Gregor Mendel

Once the agar/Petri dish, with the mycelial bridge, was fastened to the testing platform, the plants could then be placed on each island of agar. Several different types of plants were examined, such as *Cucurbita pepo* (Yellow Crookneck Squash from Plantation Products LLC), *Cucumis sativus* (Muncher Cucumber from Plantation Products LLC), *Pisum sativum* (Alaska Pea and Cascadia Sugar Snap Pea from Plantation Products LLC), *Arabidopsis thaliana* (125680 Vial # CS 70000), and *Helianthus giganteus* (Mammoth Sunflower from Plantation Products LLC), but the selection was narrowed down to pea and cucumber plants for ease of testing.

Three methods for growing plants were tested. The first method involved growing plants in soil. Each seed was grown in an individual growing cell with dimensions 6 cm x 4 cm x 7 cm. The seeds were allowed to grow for up to 3 weeks under the AgroFlex 44 grow light system made by Sunlight Supply Inc. This system implemented four T5 fluorescent 120-Volt lights. The plant was then removed from the growing tray and the soil was washed away from the roots with gently running room temperature water until enough roots were isolated from the soil and could be placed onto the agar. This method is less desirable than the following as this likely causes the plant stress.

The second method involved growing the plants in water without any soil. This process worked best with the two cultivars of *Pisum sativum* used: Alaska Pea and Cascadia Sugar Snap Pea. The peas were first soaked in tap water for 24 hours. The peas were then placed in a damp paper towel, and then in a plastic container to trap the moisture and create high humidity conditions. The peas began to sprout anywhere from 24-48 hours after being placed in the humidity chamber. The peas were then taken out of the humidity chamber and placed under a damp paper towel in an agar dish under the grow light. Each agar dish was raised on one side to give the peas a sense of gravity for roots to follow downward and shoots to rise toward the light. After 3-5 days, the peas would begin establishing roots and shoots (Figure 6A). The plants were allowed to grow for approximately 14 days before testing (Figure 6B).

**Figure 6.**
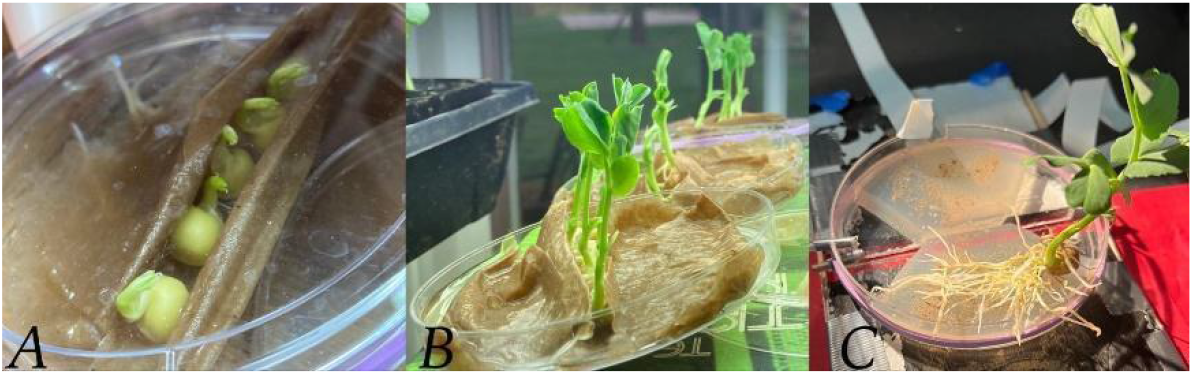
Pea Growing Process. This figure depicts the process of growing pea plants from sprouting to testing. Figure 4A depicts the pea plants just beginning to sprout in a moist paper towel enclosure. Figure 4B depicts the same pea plants further along in the growth cycle with the above-mentioned angled setup. At this stage the pea plants are ready for testing. Figure 4C depicts the pea plant removed from the moist paper towel with the roots placed on one of the agar islands.

The third method placed the plants directly on the agar, whether as seedlings or as young plants with established root systems. When grown this way, the seedlings and roots were inoculated immediately after being placed on the agar (Figures 4A and 4C). The seeds were placed in a bag filled with inoculant and shaken until thoroughly coated. For the peas with developed root systems, .05 grams of inoculant was powdered over the roots. The plants were then covered with a damp cloth to ensure the roots did not dry out (Figures 4B and 4D). The lids of the dishes were put back on to maintain moisture. For the larger pea plants, small holes were cut in the lid to allow the pea shoots to grow up and out of the dish.

The pea plants grown without soil were better suited for these experiments as no soil needed to be removed from the root system before testing. Furthermore, for the plants grown in soil, only some of the plant’s roots could be placed on the agar. Comparatively, the root systems grown in water were able to be placed in their entirety on the agar islands (Figure 6C). This greatly increased the root surface area in contact with the agar, which correspondingly improved the strength of signals detected. All roots were pulled away from the edge of the island to ensure no roots crossed the agar gap themselves, which would have created an alternate pathway for signal conduction. A moist paper towel was placed firmly over the roots to hold them in place away from the edge of the island, to ensure no root movement, and to keep the roots moist. Similar to the roots, the paper towel needed to be kept away from the agar gap as this material could transmit a signal as well. With the roots of two plants securely placed on each island of agar respectively, the testing of the mycelial bridges could now take place.

### 2.4. Mycelium Mediated Plant Electrophysiological Communication

Measuring electrical responses within the stems of plants was performed by inserting glass microelectrodes (catalog # 30-31-0 from FHC, Brunswick, ME, 04011, USA) with tips broken to jagged openings in the range of 10 to 20 μM diameter. The electrode was filled with 0.3 M KCl. The ground was placed at the base of the taproot and a small amount of silver paste made by Bare Conductive (bareconductive.com) was used to increase the conductivity between the taproot and ground wire. The electrical signals were obtained with an amplifier (Neuroprobe amplifier, A-M systems; obtained from ADInstruments, Colorado Springs, CO. 80906 USA) Recordings were performed at an acquisition rate of 20 kHz. Events were observed and analyzed with software Lab-Chart 7.0 or 8.0. The silver wires of the recording, stimulating, and ground wires were coated with chloride by soaking the wires in bleach for about 20 minutes to obtain the Ag-Cl coating. All wires were rinsed thoroughly with water prior to being used. Glass electrodes were placed within the stems with a micromanipulator under a dissecting microscope. The electrodes were inserted 1 to 2 mm into the stem of the plants. The recording setup was performed within a grounded Faraday cage and on an air table to control background noises and vibrations. Each test followed a similar protocol. This involved an agar or soil touch with fine scissors depending on which plant growing method was being used, a leaf nudge, a leaf snip, a bridge break, and another leaf snip. Each test provided different methods for examining and troubleshooting the various components of the bio-circuitry to determine if the experimental setup was properly functioning. Another function of these tests was to control for slight mechanical disturbances that could occur during the cutting of leaf. These controls helped to distinguish between movement artifacts and wound signals traveling across the agar bridge. The methods described can be illustrated with Figure 7.

**Figure 7.**
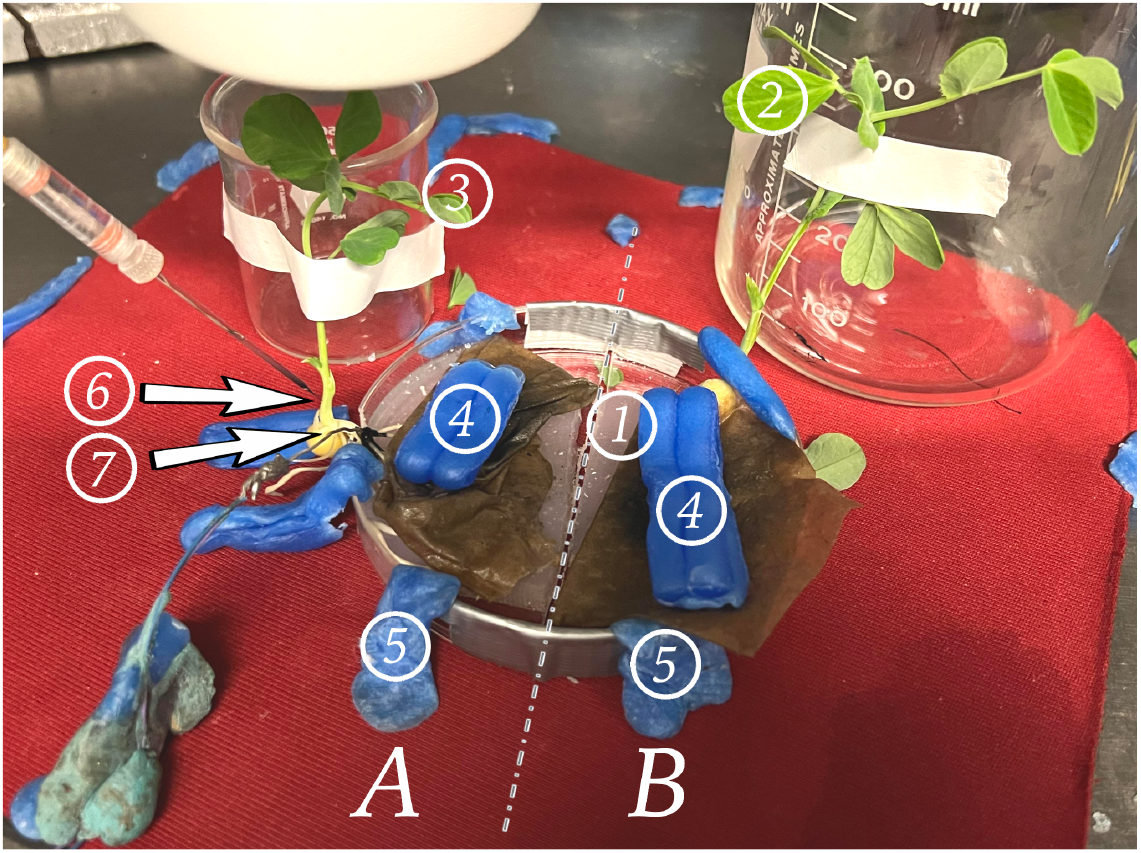
We denote the side of the dish with the glass electrode and ground wire (6 and 7) as “Side A.” The plant and agar island on the side of the setup not being recorded is denoted “Side B.” The demarcation between side A and B is denoted with the dashed line - representing the gap between the two islands of agar. Each plant is fixed in position with tape and a beaker for stem support. Wax is used to secure the dish to the testing surface which rests on an air table. The numbers indicate where different controls are performed:.

1. A patch of agar suitable for performing an agar touch on the side B.
2. A leaf suitable for a leaf nudge and leaf snip on Side B.
3. A leaf suitable for a leaf nudge and leaf snip on Side A.
4. Surgical wax is used as weights to hold down the roots under the paper towel, and to keep the roots in good contact with the agar they rest on.
5. Surgical wax is used to secure the setup to the testing pad.
6. Glass electrode inserted into the plant stem on Side A.
7. Reference electrode adhered with silver paste to the taproot of the plant on Side A.

The agar/soil touch involved inserting a pair of metal scissors into whichever medium was being used. Since both the agar and moist soil are conductive materials, signals can pass through them easily. If the scissors were inserted into the soil/agar on Side A, a large depolarization would occur, even when the individual holding the scissors was properly grounded (Figure 8A). This is due to the natural field potentials present within the human body. This agar/soil touch was implemented on Side B (Figure 8B) to determine if signals would pass across the mycelial bridge. The magnitude of the signal received corresponded to the number and density of the mycelial bridges growing between the two agar islands. A strong indication the bridges were not adequately viable/connected was when a signal was not detected during the agar/soil touch on Side B. This inadequacy could not be pinpointed, but likely was due to the dish containing very few mycelial bridges, or potentially dried out bridgings.

**Figure 8.**
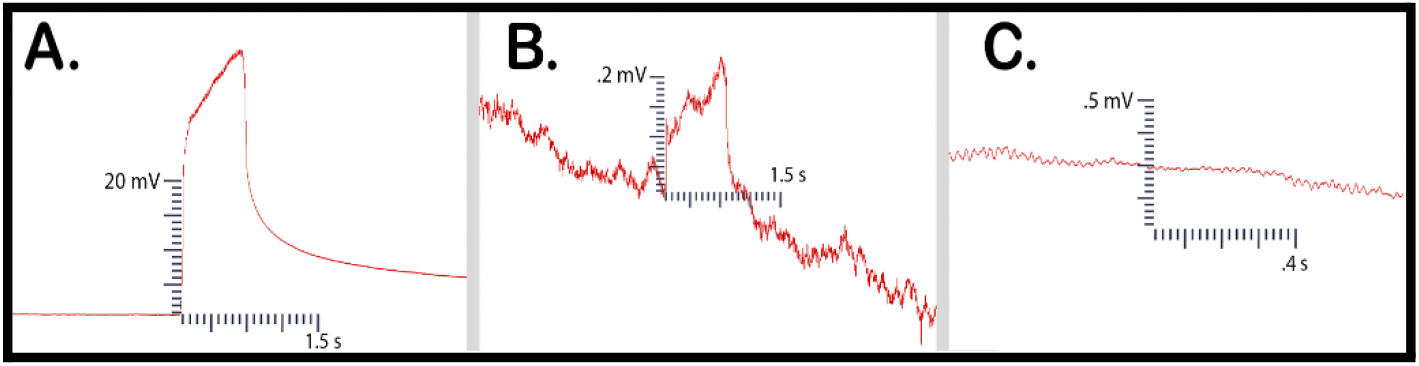
Control Examples. Figure 8A illustrates an agar touch performed on Side A. Figure 8B provides an example of an agar touch on Side B. Figure 8C is an example of not receiving a signal after a leaf bend.

If no electric signal was detected during the agar/soil touch on Side B, a moist suture thread was then placed across the two islands of agar to mimic the mycelium’s conductive role. The agar/soil was then touched once more to see if signals were detected in the recording electrode. With the suture string in place, signals were reliably and consecutively recorded. Examples of these electric potentials can be seen in Figure 13.

The next test performed was a leaf nudge. Leaf nudges help to control for artifacts in the data which can arise from plant movement. If, when nudging a leaf, a small response is detected, the plant needed to be fastened more securely into position. This involved the adhering of surgical wax around the plant stem to a stationary object. This process was repeated until no movement artifact was discernible from the background noise in the recording electrode (Figure 8C). This protocol ensured that when the leaf was snipped, responses would not be due to movement or vibration.

Leaf snips followed the leaf nudges. A leaf snip was first performed on Side B. If a signal was detected, the mycelial bridge was cut with a scalpel and another leaf snip was performed to test if a signal was still being conducted. Finally, a leaf snip was performed on the same plant being recorded in to ensure the equipment was recording properly and that a leaf snip would induce a response in the plant being monitored.

See Supplementary Materials for informative videos regarding methods and testing.

## 3. Results

The results of four trials, using *Pisum sativum* and *Cucumis sativus* in soil, and soil-less setups, are reported below. The controls used to test all setups remained the same in the four trials - implementing the soil touch, leaf nudge, leaf snip, and bridge break respectively.

### 3.1. Cucumber Tests - two trials with plants placed on inoculated agar

The first two trials included cucumbers grown in soil. Both trials consisted of three setups for a total of six setups with plants in soil. All six setups tested cucumber plants specifically. The bridges used in all six trials were selected from a larger subset of bridges and only the best bridges were selected based on the number and density of bridgings.

In trial one, following all preliminary controls mentioned in the methods, setups 1-1, 1-2, and 1-3 all achieved one or more signal conductions across the mycelial bridges from Side B to Side A. Furthermore, after the bridges were cut with a scalpel, a signal could not be transmitted from Side B to Side A. The plant signals conducted across the mycelial bridges can be seen in Figure 9.

**Figure 9.**
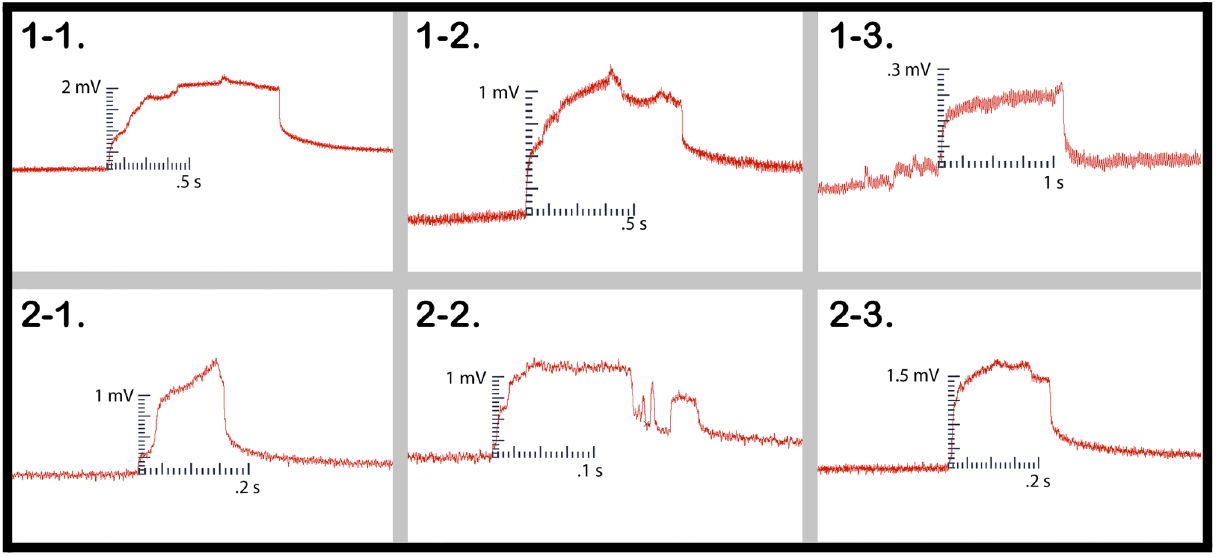
Wound Responses conducted across the bridge for Cucumber Trials 1 and 2. This figure depicts the electrical responses that were conducted from one cucumber plant to another via rhyzoelectric pathways for the six setups. The scale values on the x-axis are in seconds and the scale values on the y-axis are in millivolts.

In trial two, following all preliminary controls mentioned in the methods, setups 2-1, 2-2, and 2-3 achieved one or more signal conductions across the mycelial bridges from Side B to Side A. Furthermore, after the bridges were cut with a scalpel, a signal could not be transmitted from Side B to Side A. The plant signals conducted across the mycelial bridges can be seen in Figure 9.

The magnitude and duration of the signal likely varied based on a number of factors such as plant and root health, healthy root surface area in contact with agar, soil moisture, soil composition, and mycelial bridge health and density. The change in electrical potential and durations are listed in Table 1.

**Table 1.** This table depicts the magnitude and duration of the wound responses conducted from Side B to Side A for Cucumber trials 1 and 2.

| Cucumber Trials 1 and 2 |  |  |
| --- | --- | --- |
| Setup # | Magnitude (mV) | Duration (s) |
| 1-1 | 2.1 | 1.02 |
| 1-2 | 1 | 0.730 |
| 1-3 | 0.23 | 1.1 |
| 2-1 | 1.4 | 0.178 |
| 2-2 | 1.1 | 0.218 |
| 2-3 | 1.5 | 0.243 |
| **Average** | **1.222** | **0.582** |

### 3.2. Pea Tests - two trials with plants placed on inoculated agar

The next two trials consisted entirely of pea plants. The first pea trial tested five setups and the second pea trial tested seven setups. All 12 setups implemented pea plants grown in water except for one setup which implemented pea plants grown in soil. One soil setup was tested to compare between the two growing methods with plants of the same maturity and species with similar bridge density.

In the first trial one setup used pea plants grown in soil and four setups used pea plants grown in water. The 5 best bridges were selected from a larger population of bridges based on the number and density of bridgings. All 5 setups recorded leaf wound electric potentials in the recording electrode from Side B to A - except for one setup which only registered a response during a root wound response. Although a leaf wound response was not transmitted, a wound response in general was induced and the corresponding electric potential traveled across the mycelial bridge Figure 10. The magnitude, duration, and average values for the five setups in trial one are presented in Table 2.

**Figure 10.**
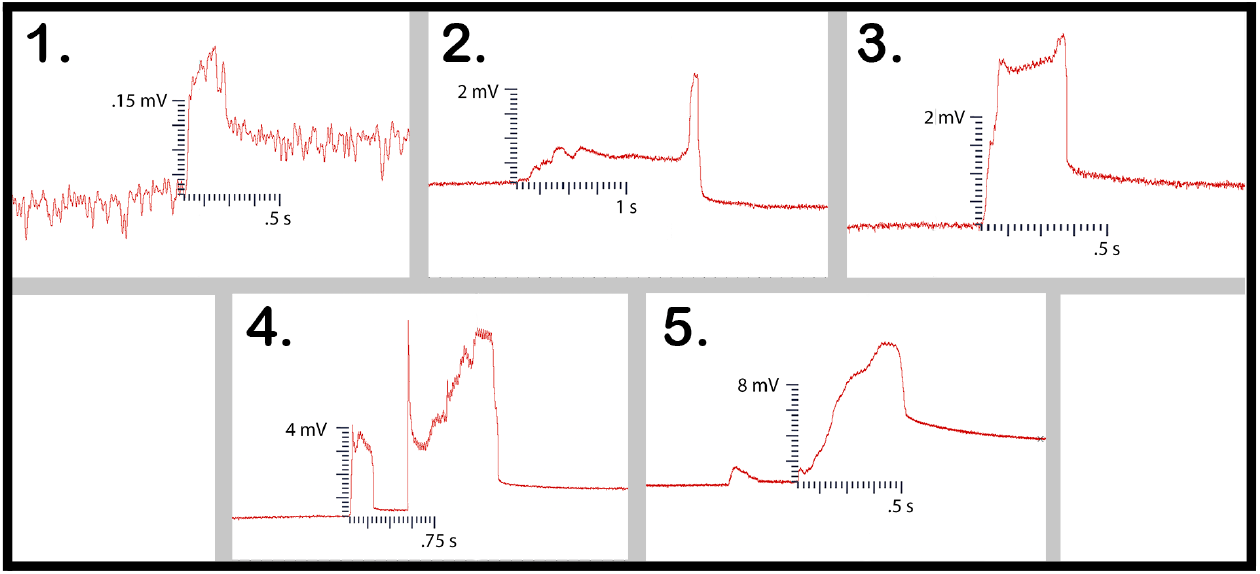
This figure depicts the 5 electrical responses that were recorded from one pea plant to another via rhyzoelectric pathways. The scale values on the x-axis are in seconds and the scale values on the y-axis are in millivolts. In setup 4, the change in electric potential and duration of the smaller response was used in the table as this was the initial response.

**Table 2.** This table depicts the magnitude and duration of the wound responses conducted from Side B to Side A for Pea Trial 1.

| Pea Trial 1 |  |  |
| --- | --- | --- |
| Setup # | Magnitude (mV) | Duration (s) |
| 1 | 0.2 | 0.217 |
| 2 | 0.6 | 1.62 |
| 3 | 3.1 | 0.348 |
| 4 | 3.4 | 0.209 |
| 5 | 9.5 | 0.529 |
| **Average** | **3.36** | **0.5846** |

Setup 1 implemented plants in soil and produced the smallest magnitude response. Setup 2 was the outlier which only responded to root snip wounding and the duration of response was longer than the other setups in the trial. Setups 4 and 5 produced small initial responses immediately followed by larger responses.

The second pea trial used plants grown in water with a moist paper towel. Seven setups were tested in this trial. The number of bridges used in the trial did not allow for a selection process in which only the best bridges were selected. Therefore, dishes that showed any bridgings in the slightest were needed due to the larger trial population. Of the seven setups which were tested, four trials were successful in conducting the wound response across the bridge to the recording electrode. After the responses were detected, the bridges were then cut with a scalpel and no further responses could be detected. These responses can be seen in trials 1, 2, 4, and 6 (Figure 11). The magnitude, duration, and average values for the four setups in trial one are shown in Table 3:

**Figure 11.**
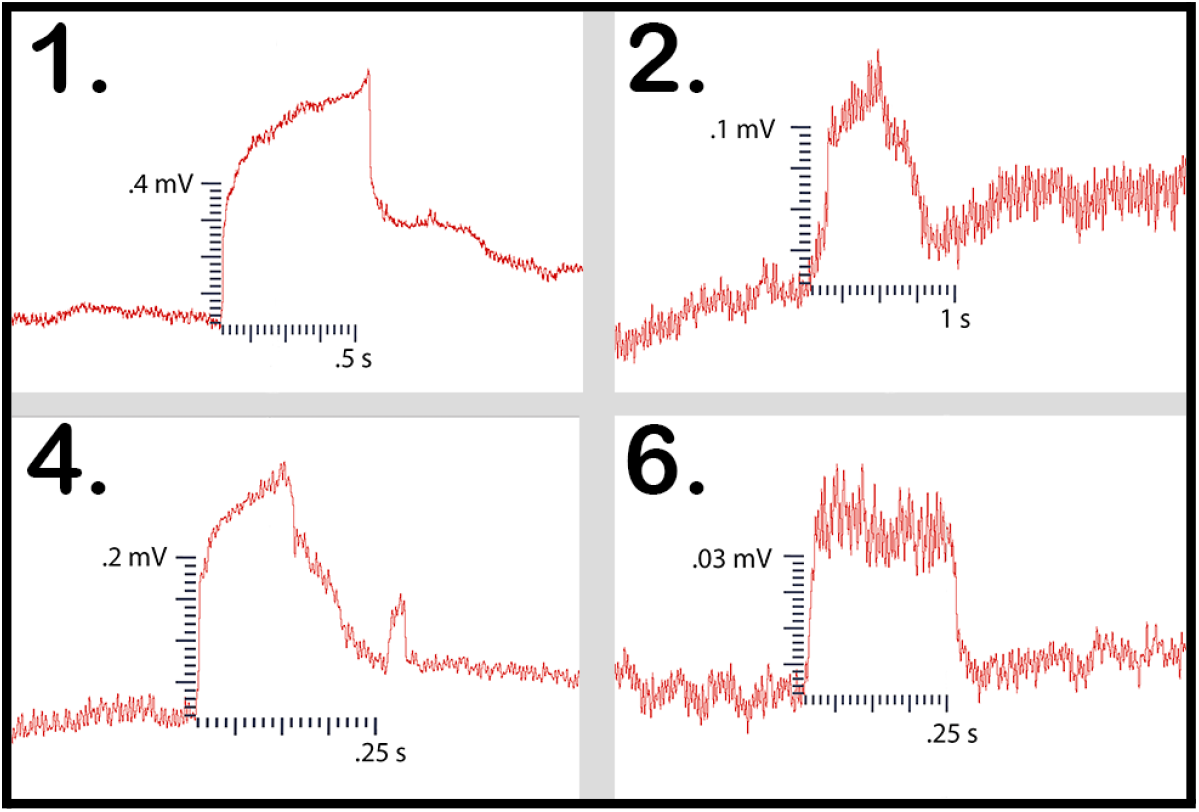
This figure depicts the electrical responses from the four trials in which a successful signal conduction occurred from one plant to another. The scale values on the x-axis are in seconds and the scale values on the y-axis are in millivolts.

**Table 3.**
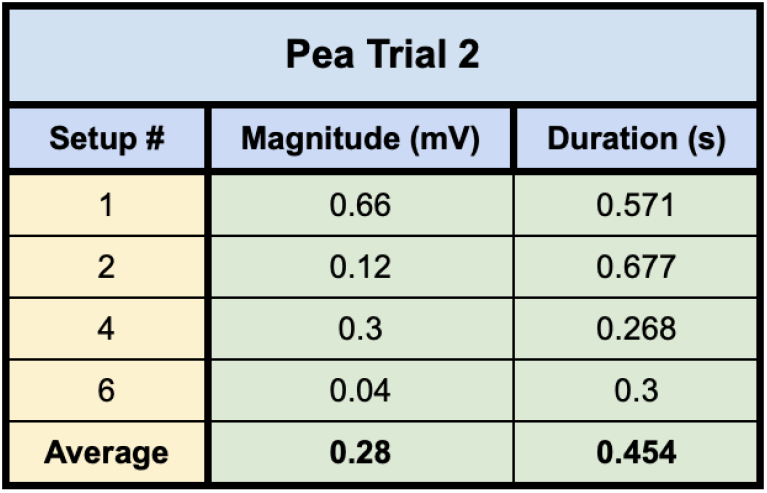
This table depicts the magnitude and duration of the wound responses conducted from Side B to Side A for Pea Trial 2.

### 3.3 Pea Tests −1 trial with plants directly inoculated and grown on agar

This trial tested seven setups with plants which had all been directly inoculated with the MycoGrow: Micronized endo/ectomycorrhizal fungi approximately two weeks before testing. Of the seven setups tested, six involved pea plants grown in tap water which had well established root systems at the time of inoculation. One setup involved direct inoculation of the pea seedlings on agar. These seedlings were then allowed to grow until they were determined to be large enough for testing. All setups were tested approximately two weeks after direct inoculation of the roots or seeds. At the time of testing, all setups were examined under a microscope prior to testing, and visual confirmation of bridges were acquired for all setups. All seven of the setups successfully conducted electric signals across their respective mycelial bridges. Upon cutting the bridges, the signals could not be conducted from the plant on Side B to the recording electrode in the plant on Side A, with the exception of one setup. Trial 3 still conducted the signal from one plant to another even after the bridge had been completely cut. Upon inspection, the authors found a likely cause for this conduction. The leaves of the pea plant on Side A (recording electrode), were slightly touching the metal table the setup was resting on, which the author was also touching with their left hand. When the author snipped the leaf of the pea plant on Side B of the setup with metal scissors, the signal likely traveled from the plant to the scissors, from the scissors to the right hand of the author, though the body of the author to their left hand which was in contact with the table, and through the table to the small portion of the pea plant in contact with the table on Side A of the setup.

### 3.4. “Sham” Bridge Conduction

Setups 3, 5, and 7 which did not successfully conduct a leaf wound response across the mycelial bridge were subjected to one final test. A moist suture thread was placed across the agar gap and the controls were repeated. This thread acted in place of the mycelial bridge, when the bridge’s health was not adequate for signal conduction. With the thread in place, an agar touch was performed on Side B. As mentioned in the methods, agar touches with metal scissors are performed to induce small electric pulses in either island of agar. Agar touches on Side B were successfully conducted across the moist suture bridge to the recording electrode in the plant on Side A for setups 3, 5, and 7 (Figure 13). The moist suture string was then removed and a signal could not be conducted across the agar gap for trials 3, 5, and 7.

**Figure 12.**
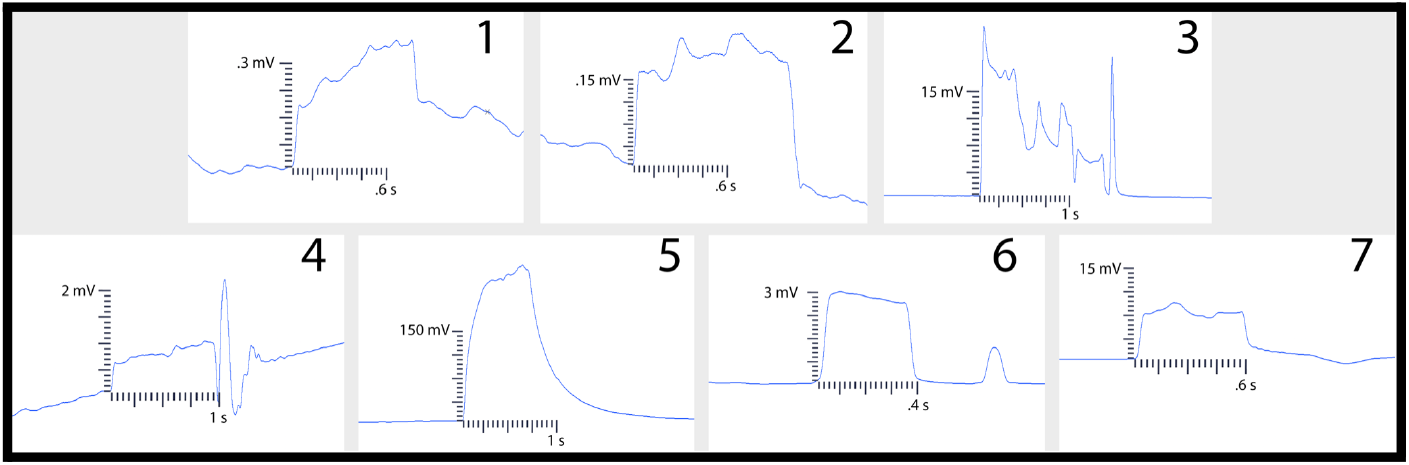
This figure depicts the electrical responses from seven trials which all produced successful signal conductions from one plant to another. The scale values on the x-axis are in seconds and the scale values on the y-axis are in millivolts.

**Figure 13.**
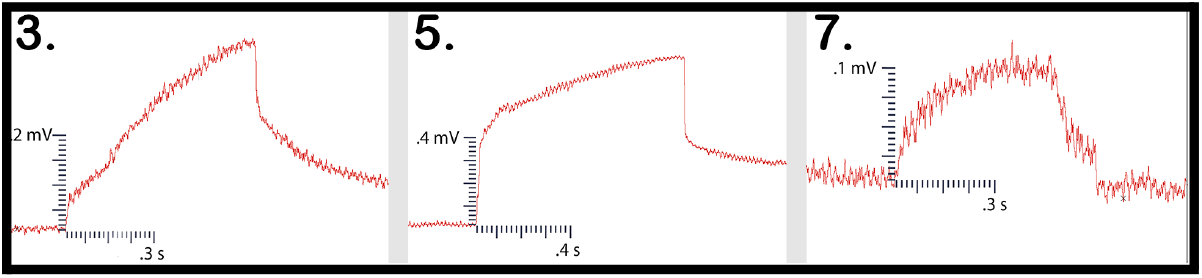
Suture Thread Electric Potentials. This figure depicts the three responses conducted from one plant to another with the moist suture thread bridging the gap.

### 3.5. Side A leaf wound response

All tests ended with a leaf snip on Side A to ensure all recording equipment was still functioning properly. The magnitudes and durations of these signals from the cucumber trials can be seen in Table 5.

**Table 4.** This table depicts the magnitude and duration of the wound responses conducted from Side B to Side A for Pea Trial 3. For complex responses, the area under the curve was taken to average the magnitude of the response.

| Pea Trial 3 |  |  |
| --- | --- | --- |
| Setup # | Magnitude (mV) | Duration (s) |
| 1 | 0.4 | 0.8 |
| 2 | 0.2 | 1.07 |
| 3 | 10.1 | 1.57 |
| 4 | 0.8 | 1.38 |
| 5 | 243 | 1.76 |
| 6 | 7.5 | 0.62 |
| 7 | 2.9 | 0.78 |
| Average | 37.8 | 1.14 |

**Table 5.** This table depicts the magnitude and duration from wound responses on Side A for Cucumber Trials 1 and 2.

| Cucumber Trials 1 and 2 |  |  |
| --- | --- | --- |
| Setup # | Magnitude (mV) | Duration (s) |
| 1-1 | 64.6 | 0.675 |
| 1-2 | 73.9 | 0.670 |
| 1-3 | 88.9 | 0.635 |
| 2-1 | 57.5 | 0.193 |
| 2-2 | 49.5 | 0.238 |
| 2-3 | 31.6 | 0.289 |
| **Average** | **61** | **0.45** |

Of the 13 pea plant setups in which the plants were placed on inoculated potato dextrose agar, Side A leaf wound responses were successfully recorded for 9 of the trials. Occasionally, the response would saturate the recording window and the true magnitude could not be determined. The magnitude and duration for the Side A wound responses obtained in the pea plant trials are presented (Table 6).

**Table 6.** This table depicts the magnitude and duration from wound responses on Side A for Pea Trials 1 and 2.

| Pea Plant Trials 1 and 2 |  |  |
| --- | --- | --- |
| Setup # | Magnitude (mV) | Duration (s) |
| 1-1 | 79.9 | 0.103 |
| 1-2 | N/A | N/A |
| 1-3 | 133 | 0.25 |
| 1-4 | 73.5 | 0.194 |
| 1-5 | 144.6 | 0.61 |
| 2-1 | N/A | N/A |
| 2-2 | 31.3 | 0.963 |
| 2-3 | N/A | N/A |
| 2-4 | 21.7 | 0.078 |
| 2-5 | 37.8 | 0.067 |
| 2-6 | 56 | 0.177 |
| 2-7 | N/A | N/A |
| **Average** | **72.225** | **0.305** |

Of the seven pea plant setups in which the plants were inoculated with the Endo/Ecto Mycorrhizal Blend, Side A leaf wound responses were successfully recorded for all seven setups. The magnitude and duration of these signals can be seen in Table 7.

**Table 7.**
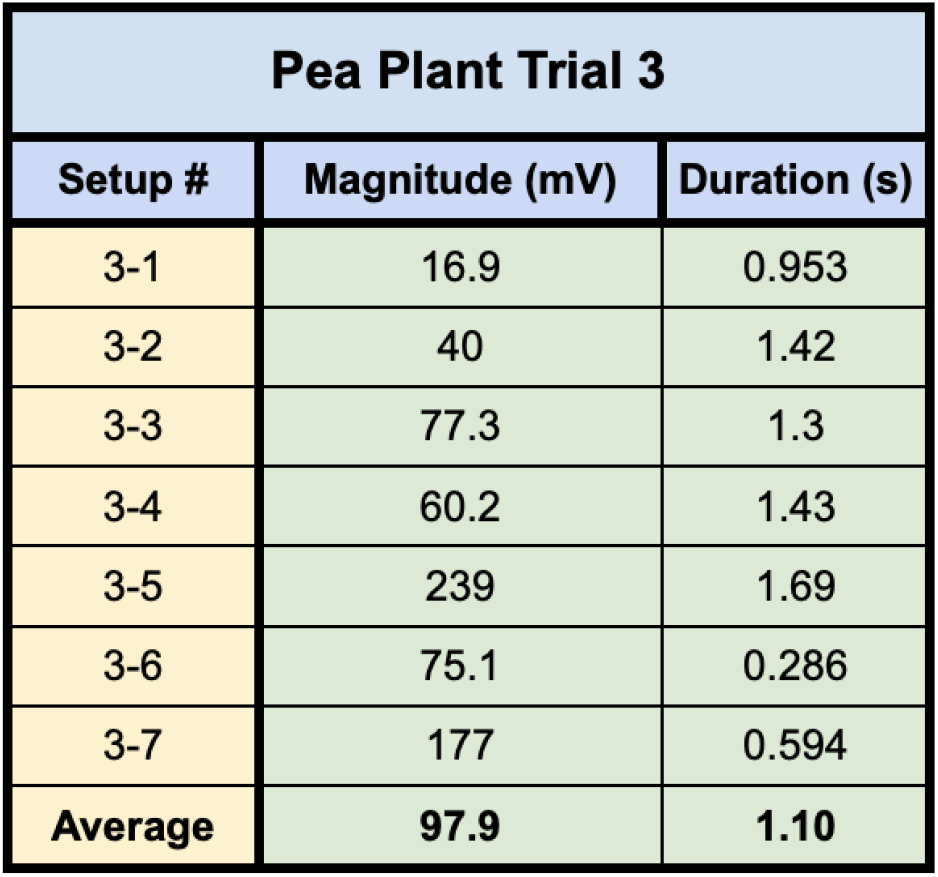
This table depicts the magnitude and duration from wound responses on Side A for Pea Trial 3.

| Pea Plant Trial 3 |  |  |
| --- | --- | --- |
| Setup # | Magnitude (mV) | Duration (s) |
| 3-1 | 16.9 | 0.953 |
| 3-2 | 40 | 1.42 |
| 3-3 | 77.3 | 1.3 |
| 3-4 | 60.2 | 1.43 |
| 3-5 | 239 | 1.69 |
| 3-6 | 75.1 | 0.286 |
| 3-7 | 177 | 0.594 |
| **Average** | **97.9** | **1.10** |

### 3.6. Signal Dampening

In comparing the average electric potential wound responses on Side A with the transmitted plant wound responses on Side B, a rough estimate of how much signal dampening, or loss, over the total length of the setup can be determined. The total distance the signal travels can be expressed using Equation 1 below.

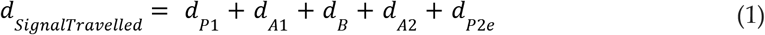

where *d*_*P*1_ is the distance from the wound site on the leaf of plant 1 to the tip of the root closest to the bridge of plant 1, *d*_*A*1_ is the distance across the agar from the root tip of plant 1 to the bridge itself, *d_B_* is the distance across the bridge spanned by the mycelium, *d*_*A*2_ is the distance across the agar from the bridge to the nearest root tip of plant 2, *d*_*P*2*e*_ is the distance from the nearest root tip of plant 2 to the recording electrode in plant 1.

For cucumbers, the average Side A wound response was 61 +/- 8.1 mV, and the average recording from the wound responses conducted from Side B was 1.22 +/- .254 mV on average. The signal received by the cucumbers on Side A is approximately 2% of the induced signal on Side B.

For Pea Trials 1 and 2, the average Side A wound response was 72.23 +/- 16.19 mV and the recording from the conducted wound response on Side B was 2 +/-1.03 mV on average. The signal received by the pea plant on Side A is approximately 2.77% of the induced signal on Side B. One likely reason for greater signal conductibility with the peas is the bare-root growing method used for testing.

For Pea Trial 3, the average Side A wound response was 97.9 +/- 28.01 and the average recording from the conducted wound response on Side B was 37.8 +/- 31.68. The reason for such a large margin of error is due to the anomaly which occurred in Setup 5 (Table 6, Setup 3-5). The signal received by the pea plant on Side A is approximately 38.6% of the induced signal on Side B for Trial 3. If we exclude the anomalous Setup 5, the signal received by the pea plant on Side A is approximately 4.9% of the induced signal on Side B. One possible reason for greater signal conduction in Pea Plant Trial 3 could be due to the method of inoculation used in Pea Trial 3.

## 4. Discussion

The methods and protocols presented provide an approach to study how plants can transmit wound induced electric potentials from one plant to another via conductive mycelial bridges. As a proof of concept, experimental setups using two species of plants were tested. Both species produced wound induced electric potentials which were successfully detected in the adjacent plant. Two methods of inoculation were tested, and both produced mycelial bridges. ITS sequencing of these fungal bridges was not performed, and so the species of the fungi cannot be definitively determined. Rather, this paper aimed to open the door for future study. Further studies could aim to compare the conductive capabilities of different strains of fungi, but this would be beyond the scope of this preliminary paper. Although contamination cannot be ruled out with either inoculation method, the results still support *a* hypothesis that hyphal filamentous threads are capable of conducting plant signals from one plant to another. The bio-circuit design isolated the conductive pathway so signal transmission could only occur across the mycelial bridge. Therefore, these responses indicate that mycelial bridges are capable of propagating plant electrophysiological signals in response to mechanical stimuli as long as the roots are alive and networked.

Theoretically, mycelium need not be within the root to pick up a plant’s electrophysiological activity. The research shown in this study indicates that fungal hyphae in contact with the root surface, or in close contact with the medium the root is in/on (such as soil or agar), is enough for these signals to be conducted from rhizome to mycelium. Therefore, ectomycorrhizal structures like Hartig’s Net should also be considered as possible rhyzoelectric pathways. This is possible so long as the medium in contact with the plant root and fungi is a good conductor of electricity, and the distance between the plant root and mycelium is such that the signal does not dissipate beyond a certain level.

Other possible explanations for how the signal was conducted from one plant to another imply the conductive pathway was not truly isolated to the mycelial bridge, but rather through another pathway. However, if preparations with few and very thin bridges were used, the success rate decreased. With inadequately healthy bridges no signal could be transmitted. Only once a wetted thread was placed across the agar gap could the signal be restored. Furthermore, setups were tested following the same experimental design, but with no mycelium present, and signals could not be conducted from one side to the other. Interestingly, the authors hypothesized that another possible avenue of conduction was potentially found for Pea Trial 3, Setup 3 (See “3.3 Pea Tests..”), but this was an exception.

These methods can be used to study the most basic plant-fungal network in vitro by implementing a type of modularity. These modules, or building blocks, consisting of plant islands and mycelial bridges can be inserted and removed to study network interactions - similar to modular components in electrical circuits. This implies larger biocircuit systems could be built with more plants, different species of plants, and more mycelial bridges.

This study can lead to addressing other questions including: how far these action potentials can disperse through a network? Can plants form chains of communication with the aid of common mycorrhizal networks? Do larger plants produce larger signals? Can mycorrhizae “eavesdrop” on these signals and respond themselves, or possibly alter the signal to change how plants respond downstream? How do the behavior of variation potentials and slow wave potentials differ or correspond to those of action potentials when looking at these local networks? Similar to neurons, are some plant responses gated - requiring the receival of action potentials from several plants, or multiple action potentials from the same plant upstream, before they themselves produce an electric potential response?

Neurons function using single and consecutive electric potential spikes [9,10]. Single and consecutive electric potential spikes also form the backbone for how *plants* detect and respond to their environment. Likewise, *fungi* produce single electric potential spikes and trains of electric potential spikes in response to conditions in their environment [11,12,13]. Fungi respond to mechanical, chemical, and optical stimulation by changing the behavior of these electric potential trains. [14,15]. This evolutionary development of sensing and responding is embedded deep within our ancient animal ancestry, and we share this common archetype for sensing, responding, and communicating with plants and fungi.

When we view an ecosystem as an incredibly complex network with underlying electrophysiological properties, we can begin applying analytic tools practiced in neurophysiology to ecology [16]. A plant’s phloem acts as a single and continuous conducting cable - likened to an axon in a single metazoan neuron [17]. For example, a neuron can be excited in the brain which triggers a cascade of neurons firing “downstream.” This is a powerful tool neurophysiologists use to establish where neuronal networks exist, without having to visually confirm which neurons are connected to each other. We could potentially apply this same tool to an ecosystem, where a large action potential is induced in a tree, and the surrounding trees and plants are recorded to see which trees receive the signal. Conclusions could be drawn from these findings as to which trees are connected underground.

Similar to the scientists aiming to create a standard model for plant signals, some researchers are going so far as to compare the electric responses of fungi to language, where different patterns of electric potential spikes represent different words. The fungi studied exhibited lexicons of up to 50 words, but the core lexicon which appeared most frequently did not exceed 15-20 words [18]. The researchers found that the distribution of the length of the spike in the spike trains corresponded to the distribution of word lengths of human languages.

A plant’s root apex, where a plant’s roots and fungi meet for the first time, generates unusually high electric fields [19,20]. Correspondingly, some fungi are known to exhibit electrotropic-like behavior - meaning they grow toward electric fields [21]. Furthermore, electric currents have been studied between roots and fungi during the formation of mycorrhizae [22]. All of these findings point to an underlying force, or central theme: networking, communication, cooperation, and trade are just as evolutionarily favorable as competition. Ecosystems strive to form networks to share information about changes in their environment because the survivability and adaptability of an organism is based on the survivability and adaptability of the ecosystem it lives in. The quicker an ecosystem can detect its ever-changing environment, the better its chances of survival and thrival.

With the insurmountable data supporting vast interconnectivity in ecosystems, we are required to view ecosystems through a different lens. A lens which looks at the ecosystem as a single organism. Similar to how we as humans are made up of a system of cells - where the cells operate independently, communicate with, combat with, and help one another, but the end result is equilibrium of the collective organism. This too is reflected in the diverse ecosystems around us in which we ourselves are but cells contributing to a network density which reaches incomputable levels of complexity.

Complexity theory, and emergence theory are wildly popular topics today with regard to many fields of study such as artificial intelligence, consciousness, psychology, ecology, network analytics, and informatics. At some resolution of the reductionist approach, the neuron is the basic building block for qualia, or subjective experience. This reductionist approach is a great start for understanding complex phenomena, but will not likely explain the phenomenological whole in its behavior beyond a certain level of complexity. This paper aims to establish the basic building block for ecological networks, but likewise, this sheds little light into how more complex ecological networks behave. A complex network does not contain just the individual nodes, but also the complex behaviors and relationships which *emerge* from this network. In other words, the Gestalt phenomena.

A study sought to determine just how interconnected these systems can be. Six 10 x10 meter plots in a Doug Fir forest were surveyed using satellite DNA marking of the 2 most common mycorrhizae in the area with respect to Xeric and Mesic soils. There were an average of 27.5 trees per plot. The number of interconnections in the plots, meaning how many trees shared genetically similar mycorrhizal symbionts, on average was 245.33 interconnections per tree pair! [23]. Lastly, we will assume mycorrhizae form symbiotic relationships with 90% of all terrestrial plants [2].

We will let τ represent the number of interconnections per tree pair. Δ will denote the number of sensory points, or nodes per tree. We will be conservative and say there are only 10,000 sensory-point-nodes per mature tree. We will then take a large population of trees as an example - the number of trees in the Amazon. This parameter will be denoted by “A.” The value “.9” simply represents that 90% of all terrestrial plants form symbiotic mycorrhizal relations. This equation is merely a model for nodal number and interconnectedness. The author notes that it is not credible to take data from Doug Fir Forest plots and apply the figures to sample sizes in the Amazon Rainforest, but this exercise is merely to provoke conversation – a drop of water on a still pond.

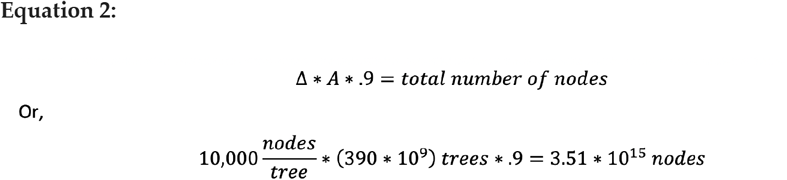

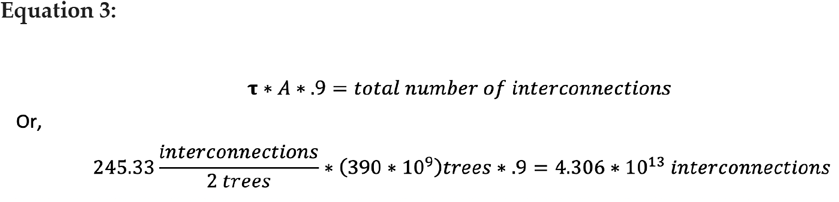

In words, 3.51 quadrillion nodes and 43.06 trillion interconnections. These figures only consider trees. Keep in mind virtually *every plant* implements electrophysiological impulses for sensing and evolution. Also keep in mind all insects, animals, and bacteria are tapping into portions of this network as well as inserting chemical and mechanical data into it. Lastly, do not forget the mycorrhizae themselves which are “so tiny that one cubic inch of soil can contain enough hyphae to stretch for 8 miles” [24], and can span continuously for kilometers [25]. If all plant life is considered part of this myco-web, the estimated values are likely orders of magnitude larger. Compare this to the brain with an estimated trillion neurons and a quadrillion synapses, or connection points. What Gestalt properties would emerge from a network larger than the human brain?

“A forest knows things. They wire themselves up underground. There are brains down there, ones our own brains aren’t shaped to see. Root plasticity, solving problems and making decisions. Fungal synapses. What else do you want to call it? Link enough trees together, and a forest grows aware.”
Richard Powers, The Overstory

## 5. Conclusion

We found that electrical signals were reliably conducted across the mycelial bridges from one plant to another upon the induction of a wound response. Our findings provide evidence that mechanical input can be communicated between plant species and opens the door to testing how this information can affect plant physiology.

## Supplementary Materials

The following supporting information can be downloaded at: www.mdpi.com/xxx/s1, Figure S1: title; Table S1: title; Video S1: title.

Plant-Fungal Electrophysiological Communication: Experimental Setup: https://www.youtube.com/watch?v=MZ6J4xLn4EU&t=0s

Plant-Fungal Electrophysiological Communication: Experimental Testing: https://www.youtube.com/watch?v=_ijgNMIOrDY&t=0s

## Author Contributions

Conceptualization, all Authors.; Data curation, all Authors; Formal analysis, all Authors.; Investigation, all Authors; Methodology, all Authors; Writing, all Authors; Writing—review and editing, all Authors. Production of supplementary materials: M.A.T. All authors have read and agreed to the published version of the manuscript.

## Funding

Department of Biology, University of Kentucky. Personal funds (R.L.C.; M.A.T.). Chellgren Endowed Funding (R.L.C.).

## Data Availability Statement

Not applicable.

## Acknowledgments

Hannah Whalen for her vigilant assistance in growing plants, Ashley Seifert for his guidance, suggestions, paper editings, and reviews, Nicholas McLetchie for suggestions and review of the paper, Joe Davis for his guidance, assistance, and inspiration, Rachel Vascessenno for her assistance with data logging, Alexandra Albakyan for her contribution to scientific progress, and Harry and Carol Thomas for their love, support, and patience..

## Conflicts of Interest

The authors declare no conflict of interest.

## Notes

https://www.youtube.com/watch?v=_ijgNMIOrDY&t=0s

https://www.youtube.com/watch?v=MZ6J4xLn4EU&t=0s

### Competing Interest Statement

The authors have declared no competing interest.

